## supplemental file 1a to 1g for "Recombination shapes the diversification of the *wtf* meiotic drivers"

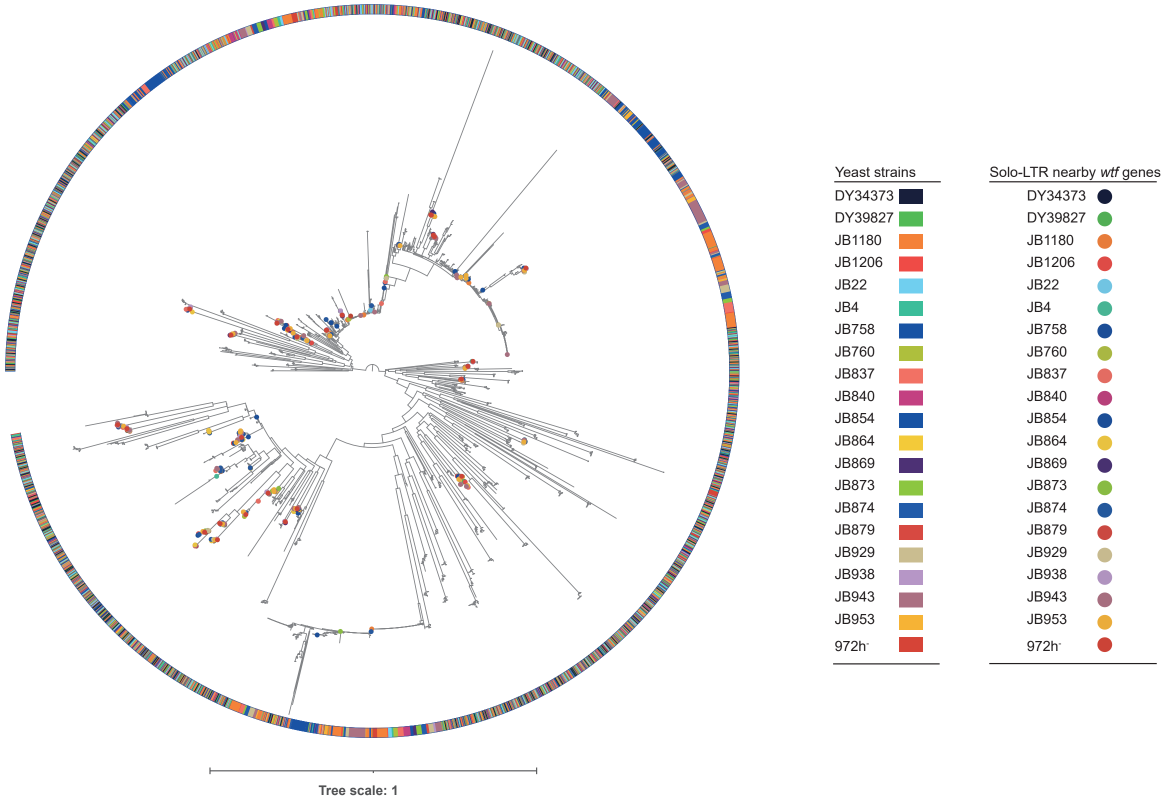


**Supplementary file 1a. The phylogenetic relationships of solo-LTRs.** The solo-LTRs nearby the *wtf* genes are labeled using solid circles in different colors indicating their source strains.

**Supplementary file 1b. Exon clusters of *wtf* genes identified in this study**

| Cluster | Exons |
|---|---|
| Cluster 1 | wtf1exon1, wtf2exon1, wtf3exon1, wtf4exon1, wtf5exon1, wtf6exon1, wtf7exon1, wtf8exon1, wtf9exon1, wtf10exon1, wtf11exon1, wtf12exon1, wtf13exon1, wtf14exon1, wtf15exon1, wtf16exon1, wtf17exon1, wtf18exon1, wtf19exon1, wtf20exon1, wtf21exon1, wtf22exon1, wtf23exon1, wtf24exon1, wtf25exon1 |
| Cluster 2 | wtf1exon3, wtf3exon1, wtf5exon3, wtf6exon3, wtf10exon3, wtf11exon3, wtf12exon3, wtf15exon3, wtf17exon3, wtf20exon3, wtf25exon3 |
| Cluster 3 | wtf4exon3, wtf8exon3, wtf9exon3, wtf13exon3, wtf16exon3, wtf18exon3, wtf19exon3, wtf21exon3, wtf22exon3, wtf23exon3, wtf24exon3 |
| Cluster 4 | wtf4exon5, wtf8exon5, wtf9exon5, wtf13exon5, wtf16exon5, wtf18exon5, wtf19exon5, wtf21exon5, wtf22exon5, wtf23exon5, wtf24exon5 |
| Cluster 5 | wtf1exon4, wtf3exon4, wtf5exon4, wtf6exon4, wtf10exon4, wtf11exon4, wtf17exon4, wtf20exon4, wtf25exon4 |
| Cluster 6 | wtf4exon2, wtf12exon2, wtf13exon2, wtf18exon2, wtf20exon2 |
| Cluster 7 | wtf4exon4, wtf8exon4, wtf9exon4, wtf13exon4, wtf16exon4, wtf18exon4, wtf19exon4, wtf21exon4, wtf22exon4, wtf23exon4, wtf24exon4 |
| Cluster 8 | wtf4exon6, wtf8exon6, wtf9exon6, wtf13exon6, wtf16exon6, wtf18exon6, wtf19exon6, wtf21exon6, wtf22exon6, wtf23exon6, wtf24exon6 |
| Cluster 9 | Wtf1exon2, wtf3exon2, wtf5exon2, wtf6exon2, wtf8exon2, wtf9exon2, wtf10exon2, wtf16exon2, wtf17exon2, wtf19exon2, wtf21exon2, wtf22exon2, wtf23exon2, wtf24exon2, wtf25exon2 |
| Cluster 10 | wtf1exon5, wtf3exon5, wtf5exon5, wtf6exon5, wtf10exon5, wtf15exon5, wtf17exon5, wtf20exon5, wtf25exon5, wtf2exon3 |

**Supplementary file 1c. *S. pombe* strains used in this study**

| **Strain** | **Assembly No.** |
|---|---|
| 972h$^-$ | GCA_000002945.2 |
| JB929 | CNA0022743 |
| JB758 | CNA0022744 |
| JB874 | CNA0022745 |
| JB943 | CNA0022746 |
| DY34373 | CNA0022747 |
| DY39827 | CNA0022748 |
| JB4 | CNA0022729 |
| JB938 | CNA0022730 |
| JB1180 | CNA0022731 |
| JB864 | CNA0022732 |
| JB22 | CNA0022733 |
| JB1206 | CNA0022734 |
| JB837 | CNA0022735 |
| JB869 | CNA0022736 |
| JB953 | CNA0022737 |
| JB760 | CNA0022738 |
| JB873 | CNA0022739 |
| JB854 | CNA0022742 |
| JB840 | CNA0022741 |
| JB879 | CNA0022740 |

**Supplementary file 1e. *wtf* genes identified in this study**

| Strain | Gene ID | Location |
|---|---|---|
| DY34373 | Segkk10arrow | 1209621-1211305 |
| DY34373 | Segkk10arrow | 1212213-1213270 |
| DY34373 | Segkk14arrow | 1038621-1039763 |
| DY34373 | Segkk14arrow | 1040827-1041973 |
| DY34373 | Segkk14arrow | 210438-211499 |
| DY34373 | Segkk14arrow | 247532-248755 |
| DY34373 | Segkk14arrow | 281656-282987 |
| DY34373 | Segkk14arrow | 318247-319601 |
| DY34373 | Segkk14arrow | 362790-364105 |
| DY34373 | Segkk14arrow | 365074-366371 |
| DY34373 | Segkk14arrow | 391303-392619 |
| DY34373 | Segkk14arrow | 411779-412435 |
| DY34373 | Segkk14arrow | 413343-414754 |
| DY34373 | Segkk14arrow | 538878-540301 |
| DY34373 | Segkk14arrow | 626636-628202 |
| DY34373 | Segkk14arrow | 629114-630164 |
| DY34373 | Segkk14arrow | 694734-696251 |
| DY34373 | Segkk14arrow | 801368-802482 |
| DY34373 | Segkk14arrow | 803588-804508 |
| DY34373 | Segkk14arrow | 853426-854855 |
| DY34373 | Segkk16arrow | 201278-202712 |
| DY34373 | Segkk16arrow | 223913-225455 |
| DY34373 | Segkk16arrow | 226370-227415 |
| DY34373 | Segkk16arrow | 280701-281803 |
| DY34373 | Segkk16arrow | 282715-284312 |
| DY34373 | Segkk16arrow | 30392-31739 |
| DY34373 | Segkk16arrow | 306710-307617 |
| DY34373 | Segkk16arrow | 413401-414892 |
| DY34373 | Segkk16arrow | 415804-417291 |
| DY34373 | Segkk16arrow | 475845-476902 |
| DY34373 | Segkk16arrow | 605589-606916 |
| DY34373 | Segkk16arrow | 607829-608878 |
| DY34373 | Segkk17arrow | 488376-489752 |
| DY34373 | Segkk17arrow | 624940-626379 |
| DY39827 | Segkk12arrow | 450420-451469 |
| DY39827 | Segkk12arrow | 452382-453709 |
| DY39827 | Segkk12arrow | 582399-583456 |
| DY39827 | Segkk12arrow | 642044-643531 |

| | | |
|---|---|---|
| DY39827 | Segkk12arrow | 644443-645934 |
| DY39827 | Segkk12arrow | 751722-752629 |
| DY39827 | Segkk12arrow | 775027-776624 |
| DY39827 | Segkk12arrow | 777536-778638 |
| DY39827 | Segkk12arrow | 831925-832970 |
| DY39827 | Segkk12arrow | 833885-835436 |
| DY39827 | Segkk12arrow | 856632-858100 |
| DY39827 | Segkk15arrow | 467374-468431 |
| DY39827 | Segkk15arrow | 469339-471009 |
| DY39827 | Segkk17arrow | 1021572-1022903 |
| DY39827 | Segkk17arrow | 1055804-1057027 |
| DY39827 | Segkk17arrow | 1093062-1094123 |
| DY39827 | Segkk17arrow | 259169-260271 |
| DY39827 | Segkk17arrow | 261053-262199 |
| DY39827 | Segkk17arrow | 263263-264405 |
| DY39827 | Segkk17arrow | 449766-451195 |
| DY39827 | Segkk17arrow | 500115-501032 |
| DY39827 | Segkk17arrow | 502141-503255 |
| DY39827 | Segkk17arrow | 608373-609891 |
| DY39827 | Segkk17arrow | 674458-675508 |
| DY39827 | Segkk17arrow | 676420-677986 |
| DY39827 | Segkk17arrow | 764319-765742 |
| DY39827 | Segkk17arrow | 889803-891213 |
| DY39827 | Segkk17arrow | 892121-892777 |
| DY39827 | Segkk17arrow | 911935-913252 |
| DY39827 | Segkk17arrow | 938186-939553 |
| DY39827 | Segkk17arrow | 940452-941767 |
| DY39827 | Segkk17arrow | 984956-986310 |
| DY39827 | Segkk20arrow | 1009389-1010828 |
| DY39827 | Segkk20arrow | 1146138-1147514 |
| JB1180 | unitig_16quiver | 197269-198324 |
| JB1180 | unitig_19quiver | 167517-168555 |
| JB1180 | unitig_19quiver | 174386-175879 |
| JB1180 | unitig_19quiver | 21250-22439 |
| JB1180 | unitig_19quiver | 310734-312373 |
| JB1180 | unitig_65quiver | 225656-226737 |
| JB1180 | unitig_65quiver | 262948-264007 |
| JB1180 | unitig_65quiver | 296894-298243 |
| JB1180 | unitig_65quiver | 333499-334874 |
| JB1180 | unitig_65quiver | 377964-379310 |
| JB1180 | unitig_65quiver | 380219-381528 |

| | | |
|---|---|---|
| JB1180 | unitig_65quiver | 411417-412890 |
| JB1180 | unitig_65quiver | 433134-434139 |
| JB1180 | unitig_65quiver | 435046-436695 |
| JB1180 | unitig_66quiver | 126845-128188 |
| JB1180 | unitig_66quiver | 240251-241368 |
| JB1180 | unitig_66quiver | 242834-243754 |
| JB1180 | unitig_66quiver | 292660-294150 |
| JB1180 | unitig_66quiver | 472818-473959 |
| JB1180 | unitig_66quiver | 475023-476168 |
| JB1180 | unitig_66quiver | 476950-478052 |
| JB1180 | unitig_66quiver | 54056-55583 |
| JB1180 | unitig_66quiver | 56374-57423 |
| JB1180 | unitig_67quiver | 480426-481475 |
| JB1180 | unitig_67quiver | 482395-483641 |
| JB1180 | unitig_74quiver | 101258-102683 |
| JB1180 | unitig_74quiver | 123788-124900 |
| JB1180 | unitig_74quiver | 177336-178438 |
| JB1180 | unitig_74quiver | 179339-180948 |
| JB1180 | unitig_9quiver | 134352-135837 |
| JB1180 | unitig_9quiver | 194401-195730 |
| JB1180 | unitig_9quiver | 196637-198086 |
| JB1180 | unitig_9quiver | 308873-309778 |
| JB1180 | unitig_9quiver | 4252-5711 |
| JB1206 | Segkk17arrow | 259077-260179 |
| JB1206 | Segkk1arrow | 3666130-3667569 |
| JB1206 | Segkk1arrow | 3802765-3804141 |
| JB1206 | Segkk2arrow | 106365-107467 |
| JB1206 | Segkk2arrow | 108379-109976 |
| JB1206 | Segkk2arrow | 132373-133280 |
| JB1206 | Segkk2arrow | 239061-240553 |
| JB1206 | Segkk2arrow | 241465-242951 |
| JB1206 | Segkk2arrow | 301533-302590 |
| JB1206 | Segkk2arrow | 431277-432604 |
| JB1206 | Segkk2arrow | 433517-434566 |
| JB1206 | Segkk2arrow | 49570-51112 |
| JB1206 | Segkk2arrow | 52027-53072 |
| JB1206 | Segkk3arrow | 227537-228971 |
| JB1206 | Segkk3arrow | 250161-251701 |
| JB1206 | Segkk3arrow | 252615-253659 |
| JB1206 | Segkk3arrow | 56653-58000 |
| JB1206 | Segkk5arrow | 1081070-1082220 |

| | | |
|---|---|---|
| JB1206 | Segkk5arrow | 1083284-1084430 |
| JB1206 | Segkk5arrow | 252991-254052 |
| JB1206 | Segkk5arrow | 290086-291309 |
| JB1206 | Segkk5arrow | 324209-325540 |
| JB1206 | Segkk5arrow | 360801-362155 |
| JB1206 | Segkk5arrow | 405332-406647 |
| JB1206 | Segkk5arrow | 407616-408913 |
| JB1206 | Segkk5arrow | 433846-435162 |
| JB1206 | Segkk5arrow | 454322-454978 |
| JB1206 | Segkk5arrow | 455886-457297 |
| JB1206 | Segkk5arrow | 4760631-4762301 |
| JB1206 | Segkk5arrow | 4763209-4764266 |
| JB1206 | Segkk5arrow | 581352-582775 |
| JB1206 | Segkk5arrow | 669101-670666 |
| JB1206 | Segkk5arrow | 671578-672628 |
| JB1206 | Segkk5arrow | 737198-738716 |
| JB1206 | Segkk5arrow | 843828-844942 |
| JB1206 | Segkk5arrow | 846128-846968 |
| JB1206 | Segkk5arrow | 895890-897319 |
| JB22 | Segkk1arrow | 1026313-1027455 |
| JB22 | Segkk1arrow | 1028509-1029606 |
| JB22 | Segkk1arrow | 208616-209680 |
| JB22 | Segkk1arrow | 246297-247597 |
| JB22 | Segkk1arrow | 281537-283034 |
| JB22 | Segkk1arrow | 318263-319613 |
| JB22 | Segkk1arrow | 362999-364327 |
| JB22 | Segkk1arrow | 407417-408573 |
| JB22 | Segkk1arrow | 409486-411057 |
| JB22 | Segkk1arrow | 620195-621725 |
| JB22 | Segkk1arrow | 622515-623563 |
| JB22 | Segkk1arrow | 688055-689365 |
| JB22 | Segkk1arrow | 794734-795851 |
| JB22 | Segkk1arrow | 797318-798238 |
| JB22 | Segkk1arrow | 846981-848552 |
| JB22 | Segkk6arrow | 395830-396914 |
| JB22 | Segkk6arrow | 523993-525345 |
| JB22 | Segkk6arrow | 583737-585053 |
| JB22 | Segkk6arrow | 690802-691708 |
| JB22 | Segkk6arrow | 714065-715183 |
| JB22 | Segkk6arrow | 767353-768467 |
| JB22 | Segkk6arrow | 789619-791100 |

| | | |
|---|---|---|
| JB22 | Segkk6arrow | 792013-793114 |
| JB22 | Segkk7arrow | 488524-489579 |
| JB22 | Segkk9arrow | 18028-18418 |
| JB4 | Segkk4arrow | 1046454-1047495 |
| JB4 | Segkk4arrow | 1182731-1183920 |
| JB4 | Segkk5arrow | 1070240-1071382 |
| JB4 | Segkk5arrow | 1072446-1073591 |
| JB4 | Segkk5arrow | 1074373-1075475 |
| JB4 | Segkk5arrow | 241165-242248 |
| JB4 | Segkk5arrow | 278509-279815 |
| JB4 | Segkk5arrow | 312667-313800 |
| JB4 | Segkk5arrow | 349051-350573 |
| JB4 | Segkk5arrow | 393810-395395 |
| JB4 | Segkk5arrow | 420373-421813 |
| JB4 | Segkk5arrow | 441952-443003 |
| JB4 | Segkk5arrow | 443909-445364 |
| JB4 | Segkk5arrow | 572651-574169 |
| JB4 | Segkk5arrow | 665487-667024 |
| JB4 | Segkk5arrow | 667937-668985 |
| JB4 | Segkk5arrow | 733497-734807 |
| JB4 | Segkk5arrow | 836974-838091 |
| JB4 | Segkk5arrow | 839558-840478 |
| JB4 | Segkk5arrow | 889396-890460 |
| JB4 | Segkk6arrow | 1246750-1247805 |
| JB4 | Segkk7arrow | 6676-7708 |
| JB4 | Segkk8arrow | 463177-464089 |
| JB4 | Segkk8arrow | 465002-466523 |
| JB4 | Segkk8arrow | 593215-594771 |
| JB4 | Segkk8arrow | 653350-654683 |
| JB4 | Segkk8arrow | 655589-657041 |
| JB4 | Segkk8arrow | 762941-763846 |
| JB4 | Segkk8arrow | 786211-787635 |
| JB4 | Segkk8arrow | 788543-789591 |
| JB4 | Segkk8arrow | 842516-843630 |
| JB4 | Segkk8arrow | 864805-866288 |
| JB4 | Segkk8arrow | 867201-868335 |
| JB758 | Segkk10arrow | 520099-521156 |
| JB758 | Segkk10arrow | 526386-528035 |
| JB758 | Segkk14arrow | 470107-471405 |
| JB758 | Segkk14arrow | 607046-608084 |
| JB758 | Segkk14arrow | 608995-610491 |

| | | |
|---|---|---|
| JB758 | Segkk17arrow | 182477-183903 |
| JB758 | Segkk17arrow | 205078-206352 |
| JB758 | Segkk17arrow | 207520-209019 |
| JB758 | Segkk17arrow | 209922-211095 |
| JB758 | Segkk17arrow | 264546-265455 |
| JB758 | Segkk17arrow | 292392-293299 |
| JB758 | Segkk17arrow | 399092-400534 |
| JB758 | Segkk17arrow | 459113-460533 |
| JB758 | Segkk17arrow | 599268-600319 |
| JB758 | Segkk18arrow | 1079840-1080982 |
| JB758 | Segkk18arrow | 1086679-1087781 |
| JB758 | Segkk18arrow | 235118-236179 |
| JB758 | Segkk18arrow | 275960-277254 |
| JB758 | Segkk18arrow | 308410-309551 |
| JB758 | Segkk18arrow | 344797-346117 |
| JB758 | Segkk18arrow | 389336-390715 |
| JB758 | Segkk18arrow | 415691-417164 |
| JB758 | Segkk18arrow | 442225-443330 |
| JB758 | Segkk18arrow | 567397-568896 |
| JB758 | Segkk18arrow | 670196-671753 |
| JB758 | Segkk18arrow | 673030-674116 |
| JB758 | Segkk18arrow | 739028-740594 |
| JB758 | Segkk18arrow | 845712-846826 |
| JB758 | Segkk18arrow | 847904-848824 |
| JB758 | Segkk18arrow | 897797-899227 |
| JB760 | Segkk19arrow | 489680-490735 |
| JB760 | Segkk22arrow | 372754-373891 |
| JB760 | Segkk23arrow | 105406-106977 |
| JB760 | Segkk23arrow | 284753-285895 |
| JB760 | Segkk23arrow | 286949-288046 |
| JB760 | Segkk23arrow | 53159-54276 |
| JB760 | Segkk23arrow | 55743-56663 |
| JB760 | Segkk27arrow | 269344-269478 |
| JB760 | Segkk27arrow | 56792-57839 |
| JB760 | Segkk27arrow | 58631-60161 |
| JB760 | Segkk36arrow | 1203-2338 |
| JB760 | Segkk36arrow | 126351-127947 |
| JB760 | Segkk36arrow | 166578-167878 |
| JB760 | Segkk36arrow | 204496-205560 |
| JB760 | Segkk36arrow | 45419-46748 |
| JB760 | Segkk36arrow | 601391-602490 |

| | | |
|---|---|---|
| JB760 | Segkk36arrow | 603404-604858 |
| JB760 | Segkk36arrow | 626002-627116 |
| JB760 | Segkk36arrow | 679297-680415 |
| JB760 | Segkk36arrow | 702774-703682 |
| JB760 | Segkk36arrow | 809608-810926 |
| JB760 | Segkk36arrow | 869329-870681 |
| JB760 | Segkk36arrow | 90157-91506 |
| JB760 | Segkk36arrow | 997762-998846 |
| JB760 | Segkk39arrow | 94304-94694 |
| JB837 | Segkk0arrow | 532254-533405 |
| JB837 | Segkk11arrow | 1370877-1371979 |
| JB837 | Segkk11arrow | 1372761-1374037 |
| JB837 | Segkk11arrow | 1599971-1600888 |
| JB837 | Segkk11arrow | 1601968-1603082 |
| JB837 | Segkk11arrow | 1705309-1706968 |
| JB837 | Segkk11arrow | 1771478-1772525 |
| JB837 | Segkk11arrow | 1773438-1775045 |
| JB837 | Segkk11arrow | 1861458-1863043 |
| JB837 | Segkk11arrow | 1992119-1993487 |
| JB837 | Segkk11arrow | 1994395-1995442 |
| JB837 | Segkk11arrow | 2015566-2017132 |
| JB837 | Segkk11arrow | 2042009-2043565 |
| JB837 | Segkk11arrow | 2044474-2045806 |
| JB837 | Segkk11arrow | 204843-206404 |
| JB837 | Segkk11arrow | 2088936-2090557 |
| JB837 | Segkk11arrow | 2125764-2127316 |
| JB837 | Segkk11arrow | 2165109-2166531 |
| JB837 | Segkk11arrow | 2202789-2203849 |
| JB837 | Segkk11arrow | 227569-228617 |
| JB837 | Segkk11arrow | 25987-27334 |
| JB837 | Segkk11arrow | 281831-282879 |
| JB837 | Segkk11arrow | 283787-285178 |
| JB837 | Segkk11arrow | 307551-308458 |
| JB837 | Segkk11arrow | 414369-415820 |
| JB837 | Segkk11arrow | 474387-475971 |
| JB837 | Segkk11arrow | 604770-606325 |
| JB837 | Segkk11arrow | 607232-608145 |
| JB837 | Segkk3arrow | 445097-446635 |
| JB837 | Segkk3arrow | 586777-587905 |
| JB840 | Segkk15arrow | 1020675-1022172 |
| JB840 | Segkk15arrow | 1055410-1056716 |

| | | |
|---|---|---|
| JB840 | Segkk15arrow | 1092978-1094061 |
| JB840 | Segkk15arrow | 259654-260756 |
| JB840 | Segkk15arrow | 261538-262685 |
| JB840 | Segkk15arrow | 263749-264891 |
| JB840 | Segkk15arrow | 444220-445284 |
| JB840 | Segkk15arrow | 499157-500074 |
| JB840 | Segkk15arrow | 501544-502661 |
| JB840 | Segkk15arrow | 604850-606193 |
| JB840 | Segkk15arrow | 670699-671746 |
| JB840 | Segkk15arrow | 763094-764766 |
| JB840 | Segkk15arrow | 888858-890506 |
| JB840 | Segkk15arrow | 891413-892548 |
| JB840 | Segkk15arrow | 912693-914244 |
| JB840 | Segkk15arrow | 939232-940645 |
| JB840 | Segkk15arrow | 984132-985448 |
| JB840 | Segkk18arrow | 474387-475442 |
| JB840 | Segkk21arrow | 503547-504836 |
| JB840 | Segkk21arrow | 640079-641117 |
| JB840 | Segkk21arrow | 642027-643523 |
| JB840 | Segkk22arrow | 185273-186699 |
| JB840 | Segkk22arrow | 262300-263208 |
| JB840 | Segkk22arrow | 285576-286481 |
| JB840 | Segkk22arrow | 392340-393792 |
| JB840 | Segkk22arrow | 394698-396031 |
| JB840 | Segkk22arrow | 454619-456105 |
| JB840 | Segkk22arrow | 5521-6867 |
| JB840 | Segkk22arrow | 584805-585865 |
| JB854 | Segkk0arrow | 1239062-1240117 |
| JB854 | Segkk1arrow | 259955-261057 |
| JB854 | Segkk1arrow | 261839-262986 |
| JB854 | Segkk1arrow | 264050-265192 |
| JB854 | Segkk20arrow | 1025630-1027126 |
| JB854 | Segkk20arrow | 1028036-1029074 |
| JB854 | Segkk20arrow | 1164368-1165557 |
| JB854 | Segkk21arrow | 186363-187789 |
| JB854 | Segkk21arrow | 208959-210073 |
| JB854 | Segkk21arrow | 262263-263378 |
| JB854 | Segkk21arrow | 285717-286622 |
| JB854 | Segkk21arrow | 392476-393937 |
| JB854 | Segkk21arrow | 452525-454011 |
| JB854 | Segkk21arrow | 582661-583743 |

| | | |
|---|---|---|
| JB854 | Segkk21arrow | 6763-8109 |
| JB854 | Segkk6arrow | 221188-222271 |
| JB854 | Segkk6arrow | 258521-259580 |
| JB854 | Segkk6arrow | 292468-293819 |
| JB854 | Segkk6arrow | 329076-330598 |
| JB854 | Segkk6arrow | 373628-374956 |
| JB854 | Segkk6arrow | 399992-401543 |
| JB854 | Segkk6arrow | 421686-422821 |
| JB854 | Segkk6arrow | 423728-425376 |
| JB854 | Segkk6arrow | 549467-551139 |
| JB854 | Segkk6arrow | 642482-643529 |
| JB854 | Segkk6arrow | 708032-709375 |
| JB854 | Segkk6arrow | 815265-816382 |
| JB854 | Segkk6arrow | 817849-818769 |
| JB854 | Segkk6arrow | 867686-868750 |
| JB864 | Segkk25arrow | 329957-331016 |
| JB864 | Segkk46arrow | 258005-259055 |
| JB864 | Segkk46arrow | 259966-261501 |
| JB864 | Segkk46arrow | 390254-391822 |
| JB864 | Segkk46arrow | 450412-451766 |
| JB864 | Segkk46arrow | 557593-558497 |
| JB864 | Segkk46arrow | 580860-581768 |
| JB864 | Segkk46arrow | 635138-636242 |
| JB864 | Segkk46arrow | 657401-658828 |
| JB864 | Segkk46arrow | 836747-838092 |
| JB864 | Segkk51arrow | 152490-153592 |
| JB864 | Segkk51arrow | 154374-155521 |
| JB864 | Segkk51arrow | 156585-157727 |
| JB864 | Segkk51arrow | 330673-332197 |
| JB864 | Segkk51arrow | 381145-382062 |
| JB864 | Segkk51arrow | 383173-384287 |
| JB864 | Segkk51arrow | 490948-492292 |
| JB864 | Segkk51arrow | 556711-557760 |
| JB864 | Segkk51arrow | 558664-560176 |
| JB864 | Segkk51arrow | 646575-648260 |
| JB864 | Segkk51arrow | 772345-773996 |
| JB864 | Segkk51arrow | 774903-776035 |
| JB864 | Segkk51arrow | 796175-797767 |
| JB864 | Segkk51arrow | 822800-824216 |
| JB864 | Segkk51arrow | 867360-868737 |
| JB864 | Segkk51arrow | 903995-905346 |

| | | |
|---|---|---|
| JB864 | Segkk51arrow | 938240-939300 |
| JB864 | Segkk51arrow | 975536-976619 |
| JB864 | Segkk69arrow | 1000532-1002027 |
| JB864 | Segkk69arrow | 1002938-1003976 |
| JB864 | Segkk69arrow | 1139157-1140445 |
| JB869 | Segkk11arrow | 1064537-1065679 |
| JB869 | Segkk11arrow | 1066733-1067830 |
| JB869 | Segkk11arrow | 1832099-1833177 |
| JB869 | Segkk11arrow | 247072-248137 |
| JB869 | Segkk11arrow | 284754-286054 |
| JB869 | Segkk11arrow | 309999-321496 |
| JB869 | Segkk11arrow | 356720-358163 |
| JB869 | Segkk11arrow | 401581-402909 |
| JB869 | Segkk11arrow | 446003-447159 |
| JB869 | Segkk11arrow | 448071-449642 |
| JB869 | Segkk11arrow | 659174-660701 |
| JB869 | Segkk11arrow | 661492-662540 |
| JB869 | Segkk11arrow | 727048-728358 |
| JB869 | Segkk11arrow | 833388-834505 |
| JB869 | Segkk11arrow | 835972-836892 |
| JB869 | Segkk11arrow | 885665-887236 |
| JB869 | Segkk2arrow | 5066916-5067971 |
| JB869 | Segkk5arrow | 626143-626533 |
| JB869 | Segkk6arrow | 106763-107881 |
| JB869 | Segkk6arrow | 160051-161165 |
| JB869 | Segkk6arrow | 182316-183818 |
| JB869 | Segkk6arrow | 184731-185865 |
| JB869 | Segkk6arrow | 83493-84399 |
| JB869 | Segkk9arrow | 136386-137738 |
| JB869 | Segkk9arrow | 264911-265994 |
| JB869 | Segkk9arrow | 76666-77984 |
| JB873 | Segkk0arrow | 10735-11645 |
| JB873 | Segkk0arrow | 12521-13578 |
| JB873 | Segkk0arrow | 142759-144220 |
| JB873 | Segkk0arrow | 202804-204360 |
| JB873 | Segkk0arrow | 35952-36860 |
| JB873 | Segkk18arrow | 34019-35479 |
| JB873 | Segkk18arrow | 36406-37620 |
| JB873 | Segkk18arrow | 38796-39846 |
| JB873 | Segkk24arrow | 37367-38556 |
| JB873 | Segkk34arrow | 98215-99256 |

| | | |
|---|---|---|
| JB873 | Segkk38arrow | 6533-7960 |
| JB873 | Segkk39arrow | 201251-202334 |
| JB873 | Segkk39arrow | 244079-244632 |
| JB873 | Segkk39arrow | 277495-278628 |
| JB873 | Segkk39arrow | 318220-319726 |
| JB873 | Segkk55arrow | 1102-2753 |
| JB873 | Segkk55arrow | 126875-128393 |
| JB873 | Segkk55arrow | 219710-221247 |
| JB873 | Segkk55arrow | 222160-223208 |
| JB873 | Segkk55arrow | 287721-289031 |
| JB873 | Segkk55arrow | 399369-400486 |
| JB873 | Segkk55arrow | 401953-402873 |
| JB873 | Segkk55arrow | 451791-452855 |
| JB873 | Segkk55arrow | 632213-633354 |
| JB873 | Segkk55arrow | 634136-635238 |
| JB873 | Segkk58arrow | 72655-73710 |
| JB873 | Segkk6arrow | 41909-43496 |
| JB873 | Segkk6arrow | 68507-69946 |
| JB874 | Segkk10arrow | 1012116-1013521 |
| JB874 | Segkk10arrow | 1048816-1050107 |
| JB874 | Segkk10arrow | 1083366-1084862 |
| JB874 | Segkk10arrow | 1085773-1086885 |
| JB874 | Segkk10arrow | 1087747-1088831 |
| JB874 | Segkk10arrow | 1125393-1126457 |
| JB874 | Segkk10arrow | 281905-283007 |
| JB874 | Segkk10arrow | 283789-284934 |
| JB874 | Segkk10arrow | 285998-286916 |
| JB874 | Segkk10arrow | 463184-464612 |
| JB874 | Segkk10arrow | 517895-519415 |
| JB874 | Segkk10arrow | 520885-522002 |
| JB874 | Segkk10arrow | 617986-619329 |
| JB874 | Segkk10arrow | 688795-689843 |
| JB874 | Segkk10arrow | 690636-692163 |
| JB874 | Segkk10arrow | 778724-780396 |
| JB874 | Segkk10arrow | 904455-905752 |
| JB874 | Segkk10arrow | 906662-907764 |
| JB874 | Segkk10arrow | 937770-939279 |
| JB874 | Segkk10arrow | 967397-968929 |
| JB874 | Segkk12arrow | 451252-452302 |
| JB874 | Segkk12arrow | 453478-454692 |
| JB874 | Segkk12arrow | 455618-457078 |

| | | |
|---|---|---|
| JB874 | Segkk12arrow | 585790-587276 |
| JB874 | Segkk12arrow | 645864-647325 |
| JB874 | Segkk12arrow | 753220-754130 |
| JB874 | Segkk12arrow | 781422-782479 |
| JB874 | Segkk12arrow | 783356-784266 |
| JB874 | Segkk12arrow | 838042-838920 |
| JB874 | Segkk12arrow | 839833-841384 |
| JB874 | Segkk12arrow | 862600-864026 |
| JB874 | Segkk13arrow | 373347-374404 |
| JB874 | Segkk13arrow | 375312-376982 |
| JB874 | Segkk14arrow | 527714-528939 |
| JB874 | Segkk14arrow | 529825-531115 |
| JB874 | Segkk14arrow | 666366-667404 |
| JB874 | Segkk14arrow | 668314-669809 |
| JB879 | Segkk2arrow | 325-1375 |
| JB879 | Segkk37arrow | 27243-28340 |
| JB879 | Segkk37arrow | 29394-30536 |
| JB879 | Segkk58arrow | 42370-43680 |
| JB879 | Segkk60arrow | 251549-252604 |
| JB879 | Segkk61arrow | 185715-187033 |
| JB879 | Segkk61arrow | 2332-3446 |
| JB879 | Segkk61arrow | 245433-246785 |
| JB879 | Segkk61arrow | 55620-56738 |
| JB879 | Segkk61arrow | 79094-80000 |
| JB879 | Segkk62arrow | 223098-223488 |
| JB879 | Segkk63arrow | 156135-157235 |
| JB879 | Segkk63arrow | 158149-159678 |
| JB879 | Segkk63arrow | 372155-373726 |
| JB879 | Segkk63arrow | 374639-375795 |
| JB879 | Segkk77arrow | 111823-113393 |
| JB879 | Segkk77arrow | 162130-163047 |
| JB879 | Segkk77arrow | 164517-165634 |
| JB879 | Segkk78arrow | 117185-118682 |
| JB879 | Segkk78arrow | 152624-153924 |
| JB879 | Segkk78arrow | 190656-191586 |
| JB879 | Segkk78arrow | 35894-37222 |
| JB879 | Segkk78arrow | 80608-81957 |
| JB879 | Segkk80arrow | 50910-51994 |
| JB929 | Segkk11arrow | 2039681-2041237 |
| JB929 | Segkk11arrow | 2099818-2101150 |
| JB929 | Segkk11arrow | 2102058-2103510 |

| | | |
|---|---|---|
| JB929 | Segkk11arrow | 2214323-2215228 |
| JB929 | Segkk11arrow | 2237597-2238420 |
| JB929 | Segkk11arrow | 2291817-2292919 |
| JB929 | Segkk11arrow | 2293832-2295383 |
| JB929 | Segkk11arrow | 2316604-2318031 |
| JB929 | Segkk25arrow | 468956-470145 |
| JB929 | Segkk25arrow | 610325-611366 |
| JB929 | Segkk33arrow | 1046608-1047750 |
| JB929 | Segkk33arrow | 1048814-1049959 |
| JB929 | Segkk33arrow | 1050740-1051843 |
| JB929 | Segkk33arrow | 209581-210663 |
| JB929 | Segkk33arrow | 246916-247973 |
| JB929 | Segkk33arrow | 280838-281971 |
| JB929 | Segkk33arrow | 317222-318744 |
| JB929 | Segkk33arrow | 361961-363548 |
| JB929 | Segkk33arrow | 388560-390000 |
| JB929 | Segkk33arrow | 415933-417068 |
| JB929 | Segkk33arrow | 417975-419626 |
| JB929 | Segkk33arrow | 543657-545175 |
| JB929 | Segkk33arrow | 636485-638022 |
| JB929 | Segkk33arrow | 638935-639983 |
| JB929 | Segkk33arrow | 815311-816428 |
| JB929 | Segkk33arrow | 817895-818815 |
| JB929 | Segkk33arrow | 872655-873719 |
| JB929 | Segkk35arrow | 453092-454142 |
| JB929 | Segkk35arrow | 455318-456532 |
| JB929 | Segkk35arrow | 457458-458918 |
| JB929 | Segkk36arrow | 797191-798246 |
| JB929 | Segkk5arrow | 6814-7348 |
| JB938 | Segkk52arrow | 136303-137352 |
| JB938 | Segkk52arrow | 138282-139812 |
| JB938 | Segkk52arrow | 348986-350557 |
| JB938 | Segkk52arrow | 351470-352626 |
| JB938 | Segkk52arrow | 395708-397037 |
| JB938 | Segkk52arrow | 440434-441783 |
| JB938 | Segkk52arrow | 477016-478513 |
| JB938 | Segkk52arrow | 512469-513769 |
| JB938 | Segkk52arrow | 550392-551456 |
| JB938 | Segkk52arrow | 70475-71785 |
| JB938 | Segkk74arrow | 1031600-1032684 |
| JB938 | Segkk74arrow | 1159776-1161128 |

| | | |
|---|---|---|
| JB938 | Segkk74arrow | 1219532-1220853 |
| JB938 | Segkk74arrow | 1326790-1327696 |
| JB938 | Segkk74arrow | 1350055-1351173 |
| JB938 | Segkk74arrow | 1403345-1404459 |
| JB938 | Segkk74arrow | 1425613-1427115 |
| JB938 | Segkk74arrow | 1428028-1429129 |
| JB938 | Segkk74arrow | 263912-265054 |
| JB938 | Segkk74arrow | 266108-267205 |
| JB938 | Segkk74arrow | 32320-33437 |
| JB938 | Segkk74arrow | 34904-35824 |
| JB938 | Segkk74arrow | 84568-86139 |
| JB938 | Segkk78arrow | 229612-230667 |
| JB938 | Segkk89arrow | 1503320-1503710 |
| JB943 | Segkk0arrow | 475184-476234 |
| JB943 | Segkk0arrow | 477410-478624 |
| JB943 | Segkk0arrow | 479550-481010 |
| JB943 | Segkk0arrow | 619547-621099 |
| JB943 | Segkk0arrow | 679697-681158 |
| JB943 | Segkk0arrow | 790762-791669 |
| JB943 | Segkk0arrow | 814025-815142 |
| JB943 | Segkk0arrow | 871273-872372 |
| JB943 | Segkk0arrow | 873535-874658 |
| JB943 | Segkk0arrow | 895804-897230 |
| JB943 | Segkk29arrow | 500670-501859 |
| JB943 | Segkk29arrow | 648956-649997 |
| JB943 | Segkk8arrow | 1020057-1021579 |
| JB943 | Segkk8arrow | 1058446-1059579 |
| JB943 | Segkk8arrow | 1094621-1095927 |
| JB943 | Segkk8arrow | 1132188-1133271 |
| JB943 | Segkk8arrow | 268903-270005 |
| JB943 | Segkk8arrow | 270787-271934 |
| JB943 | Segkk8arrow | 272998-274140 |
| JB943 | Segkk8arrow | 455129-456413 |
| JB943 | Segkk8arrow | 510273-511190 |
| JB943 | Segkk8arrow | 512660-513777 |
| JB943 | Segkk8arrow | 620891-622201 |
| JB943 | Segkk8arrow | 686728-687776 |
| JB943 | Segkk8arrow | 688569-690096 |
| JB943 | Segkk8arrow | 781460-782978 |
| JB943 | Segkk8arrow | 915906-917524 |
| JB943 | Segkk8arrow | 918431-919566 |

| | | |
|---|---|---|
| JB943 | Segkk8arrow | 944111-945551 |
| JB943 | Segkk8arrow | 970504-972068 |
| JB943 | Segkk9arrow | 1251845-1253515 |
| JB943 | Segkk9arrow | 1259983-1261040 |
| JB953 | Segkk13arrow | 463525-464575 |
| JB953 | Segkk13arrow | 465751-466965 |
| JB953 | Segkk13arrow | 467892-469415 |
| JB953 | Segkk13arrow | 608071-609656 |
| JB953 | Segkk13arrow | 668238-669680 |
| JB953 | Segkk13arrow | 775435-776340 |
| JB953 | Segkk13arrow | 798732-800151 |
| JB953 | Segkk13arrow | 801050-801957 |
| JB953 | Segkk13arrow | 855406-856449 |
| JB953 | Segkk13arrow | 857367-858918 |
| JB953 | Segkk13arrow | 880138-881565 |
| JB953 | Segkk14arrow | 203260-204325 |
| JB953 | Segkk14arrow | 241251-242335 |
| JB953 | Segkk14arrow | 243197-244309 |
| JB953 | Segkk14arrow | 245220-246716 |
| JB953 | Segkk14arrow | 279975-281266 |
| JB953 | Segkk14arrow | 316561-317965 |
| JB953 | Segkk14arrow | 361156-362688 |
| JB953 | Segkk14arrow | 390772-392091 |
| JB953 | Segkk14arrow | 392971-394124 |
| JB953 | Segkk14arrow | 414292-415394 |
| JB953 | Segkk14arrow | 416305-417817 |
| JB953 | Segkk14arrow | 546848-548308 |
| JB953 | Segkk14arrow | 549216-550755 |
| JB953 | Segkk14arrow | 655872-656986 |
| JB953 | Segkk14arrow | 658092-659012 |
| JB953 | Segkk14arrow | 712931-714434 |
| JB953 | Segkk14arrow | 895233-896375 |
| JB953 | Segkk14arrow | 897151-898253 |
| JB953 | Segkk1arrow | 7943576-7945117 |
| JB953 | Segkk1arrow | 8080511-8081801 |
| JB953 | Segkk1arrow | 8082669-8083894 |
| JB953 | Segkk8arrow | 206746-207803 |
| JB953 | Segkk8arrow | 208711-210380 |

**Supplementary file 1f. Primer sequences for knockout of the *wtf* genes**

| Primer name | Sequence (5' to 3') |
|---|---|
| wtf1-U-up | GCAAGTGCTTGGAATAACTC |
| wtf1-U-dw | ACGAGGCAAGCTAAACAGATCTTTCCCCAGTGTGTTGTG |
| wtf1-D-up | GCTGTCGATTCGATACTAACGGCCTAATTATATTGCTAACTTTAC |
| wtf1-D-dw | CAAACCATCGTTTTCAATCAC |
| wtf1-Y-up | CATCCTCTTCAGTGAAAGTG |
| wtf1-Y-dw | CTTAAATTGAACACGAGGAAC |
| wtf2-U-up | AGTAAACCACTAAACCAAAGG |
| wtf2-U-dw | ACGAGGCAAGCTAAACAGATGCCTAATTATATTGTTAACTTTAC |
| wtf2-D-up | GCTGTCGATTCGATACTAACGCTTCATAATAATAAAATCATTCTTT |
| wtf2-D-dw | GCGTTGTGCGTTATGTTAAC |
| wtf2-Y-up | TGTAGGAAGAAACAAAGTTCC |
| wtf2-Y-dw | CTCGAACTGATGACGATTAG |
| wtf3-U-up | CATTTCGTCTATCTAAGTCAC |
| wtf3-U-dw | ACGAGGCAAGCTAAACAGATGAAGCCTAATTATATTGCTAAC |
| wtf3-D-up | GCTGTCGATTCGATACTAACGCTTCGTTTCATCAAGATAATTC |
| wtf3-D-dw | ATGCGAGTCTAATTATGTGTC |
| wtf3-Y-up | CTACGATGGTTAAGCATGTG |
| wtf3-Y-dw | GGATAGGAAATGCAATTGGG |
| wtf4-U-up | ACTGCGTTATTACATACTCAC |
| wtf4-U-dw | ACGAGGCAAGCTAAACAGATATCACCAACGCAGAGAAGC |
| wtf4-D-up | GCTGTCGATTCGATACTAACGGTCACTGCCTTTTTTATTCAC |
| wtf4-D-dw | CCCACCTCTACAATAATAGG |
| wtf4-Y-up | GTACTTTCCTCCTCTGAATC |
| wtf4-Y-dw | GTCTCTAGAGCATAAGACAC |
| wtf5-U-up | GGACATACTTGTAATGCTCG |
| wtf5-U-dw | ACGAGGCAAGCTAAACAGATGTCTTTTATCTTTGCTCTTTTC |
| wtf5-D-up | GCTGTCGATTCGATACTAACGTAGCTTAAGCTTGCCGAAAG |
| wtf5-D-dw | GCCTCGAATATTGCTCATAC |
| wtf5-Y-up | GATGCAAAGTAAGGTAAGTAG |
| wtf5-Y-dw | AGATCGTAATTCGTGGATTTC |
| wtf6-U-up | CCGAAGCAGGATAGATCAC |
| wtf6-U-dw | ACGAGGCAAGCTAAACAGATGCACTAATTAGGCTACACGT |
| wtf6-D-up | GCTGTCGATTCGATACTAACGCCTTTTTTATTCACCCCAACT |
| wtf6-D-dw | GGCGAGGTATAGTTGTCAC |
| wtf6-Y-up | GATTTCTGCAACGGCCAAG |
| wtf6-Y-dw | CGAAGTATCATATCAACGTAG |
| wtf7-U-up | AGAAACCGAGTTATGGTTGC |

| | |
|---|---|
| wtf7-U-dw | ACGAGGCAAGCTAAACAGATACAGGATTTCATTTCTGTTCC |
| wtf7-D-up | GCTGTCGATTCGATACTAACGTGCATAAGGAGGAAACTATAG |
| wtf7-D-dw | GAGGTAAGAATAGGGAGAAC |
| wtf7-Y-up | GACTGAGCAAGATTCATTGC |
| wtf7-Y-dw | GGAAGATAAAGTACGCATAAC |
| wtf8-U-up | CGACTATAATAACATTGCCTG |
| wtf8-U-dw | ACGAGGCAAGCTAAACAGATTCTTCATTTCATCAAGATAATTC |
| wtf8-D-up | GCTGTCGATTCGATACTAACGTGCTTTATTAATGTAGTTGTCG |
| wtf8-D-dw | ACCTAAATTGAATCCTAACGC |
| wtf8-Y-up | GCTACTGGCAATGATCGTG |
| wtf8-Y-dw | CCCATATAGAAGACATCCAG |
| wtf9-U-up | GCCTCATATACAAAAGCCAC |
| wtf9-U-dw | ACGAGGCAAGCTAAACAGATATGGCATAAAAAAAGAAGAACG |
| wtf9-D-up | GCTGTCGATTCGATACTAACGGAGCAGCATAATAGAATATTGT |
| wtf9-D-dw | GAACGAGGTTCAGTTGTAAC |
| wtf9-Y-up | AGAAGCACGTTTTAGAAGGG |
| wtf9-Y-dw | GCATCTGGTTTACCATTCTG |
| wtf10-U-up | TGCTATCAGCAATACTGCAC |
| wtf10-U-dw | ACGAGGCAAGCTAAACAGATAGTGTAATTGCTTCTGACACA |
| wtf10-D-up | GCTGTCGATTCGATACTAACGCAGGCGAAATTGGGTAGAG |
| wtf10-D-dw | CCGGTTTTAACAGATCCGG |
| wtf10-Y-up | CAGGTACTTAACCAAAGTCG |
| wtf11-U-up | CAAGCCAGTAAATACTCGAC |
| wtf11-U-dw | ACGAGGCAAGCTAAACAGATGCGTAAAAGAAAAAAAAAGTGGG |
| wtf11-D-up | GCTGTCGATTCGATACTAACGCAAACAAATTTAATGTTTACAAAATAATG |
| wtf11-D-dw | GGCAGTGAATTACTTCTGAC |
| wtf11-Y-up | TGGATTGCACACATATTGGC |
| wtf11-Y-dw | GCATTTATAGTTAGCCTGGC |
| wtf12-U-up | CGTTCCTCAGTTCAGTTATG |
| wtf12-U-dw | ACGAGGCAAGCTAAACAGATGTAATTATTCTTCATAATAATAAGATC |
| wtf12-D-up | GCTGTCGATTCGATACTAACGGCCTAATTATATTGCTAGCTTT |
| wtf12-D-dw | GCTTGGATAAAGGAATTGCC |
| wtf12-Y-up | GGCATTGACTAGCAAATTGC |
| wtf12-Y-dw | GCATAAATCGATTGCAGAGC |
| wtf13-U-up | GGCTCCTTCTTGTTGTTTAC |
| wtf13-U-dw | ACGAGGCAAGCTAAACAGATCGATGGTTTGTTATCGTAATG |
| wtf13-D-up | GCTGTCGATTCGATACTAACGGACACATTTATTCTGTCACTG |
| wtf13-D-dw | TGTATTGTATTAGAATCCGTG |
| wtf13-Y-up | GCACGTTACCACATCAAAAG |
| wtf13-Y-dw | GTCAAAATCTTCAATCTATGTG |

| | |
|---|---|
| wtf14-U-up | GAAGCGATGTAAAAACATTGG |
| wtf14-U-dw | ACGAGGCAAGCTAAACAGATCTTTTGTGAAATTATCTGCTAAG |
| wtf14-D-up | GCTGTCGATTCGATACTAACGCCTAAAAATATGTTAGAGTTTGC |
| wtf14-D-dw | GCGATTTCCTCTTACATACC |
| wtf14-Y-up | CAGCTGGCTAAGTACTTCC |
| wtf14-Y-dw | AGCTTGAGACACAGGATTAG |
| wtf15-U-up | CACACAATGATTTCTTGGTTG |
| wtf15-U-dw | ACGAGGCAAGCTAAACAGATCTGCTCCTTTTATAACGTGC |
| wtf15-D-up | GCTGTCGATTCGATACTAACGATAGCGAGCGACAAAAAAGC |
| wtf15-D-dw | GCGTTTTAGCATGTGGAGG |
| wtf15-Y-up | CCAGTCGACACCTTTAAAAC |
| wtf15-Y-dw | CCAAGCAGCATCTCAATCG |
| wtf16-U-up | AGATGAAGGAGAAATGAATGC |
| wtf16-U-dw | ACGAGGCAAGCTAAACAGATTGCTTTATTATTGTAGTTGTCG |
| wtf16-D-up | GCTGTCGATTCGATACTAACGGTTTTTGAGAAGCAAACGTTG |
| wtf16-D-dw | GCTCATAAACAACGAGAAGG |
| wtf16-Y-up | GAGCTTTACTTGCGATCTAC |
| wtf16-Y-dw | GTTAATGACGATACTCCTCC |
| wtf17-U-up | ACATTGGATAAGCATTGAAGC |
| wtf17-U-dw | ACGAGGCAAGCTAAACAGATGAAGCCTAATTATATTGCTAAC |
| wtf17-D-up | GCTGTCGATTCGATACTAACGGTTTCAAAGCAAACTGTTTTTC |
| wtf17-D-dw | CCAAGACACAAACAATAATGG |
| wtf17-Y-up | TGCAAGACTCATGTTGCGG |
| wtf17-Y-dw | CTAACCGTATCAGAATGTCC |
| wtf18-U-up | CAATGTACTTTCCTCCTCTG |
| wtf18-U-dw | ACGAGGCAAGCTAAACAGATCATTTCTCCTTCATCTTGATTC |
| wtf18-D-up | GCTGTCGATTCGATACTAACGAGGCAGTGAATTGCTTCTGA |
| wtf18-D-dw | GTGTAAGCACGACCAACAC |
| wtf18-Y-up | TCATAGAAGTCTAATGAGTTAG |
| wtf18-Y-dw | GTAGCTTCGTCAAACGCAG |
| wtf19-U-up | CGCAGTTTGGTATTGTTAGC |
| wtf19-U-dw | ACGAGGCAAGCTAAACAGATGAGGCAGTGGATTGCTTCT |
| wtf19-D-UP | GCTGTCGATTCGATACTAACGCTCCTTCATCTTGATTCTTAAT |
| wtf19-D-dw | GAGGACTTTTCAAGGAAGTG |
| wtf19-Y-up | GAAATCAATTCCGTAGCAGC |
| wtf19-Y-dw | TGTCTTCTTTCGACATTGGG |
| wtf20-U-up | GATAAAGTGCATTGAGGACC |
| wtf20-U-dw | ACGAGGCAAGCTAAACAGATCTTCTGACACATTTATTTTGTC |
| wtf20-D-up | GCTGTCGATTCGATACTAACGAGCTACGTAGTTTGGTATTCA |

| | |
|---|---|
| wtf20-D-dw | GATGCATTTATCACCCAAGG |
| wtf20-Y-up | GGTTGTGAAATTGAAGCCTC |
| wtf20-Y-dw | CTATACGTACTACGTGCTAC |
| wtf21-U-up | GCTATGAACGACCAAATGCT |
| wtf21-U-dw | ACGAGGCAAGCTAAACAGATTGAGAAGCAAAACGTTGTAGT |
| wtf21-D-up | GCTGTCGATTCGATACTAACGCTCCACGAAAAAGGTCGAG |
| wtf21-D-dw | TGCAAGTGCAATACAATCCC |
| wtf21-Y-up | CTCCACTTTATAGCACAGTG |
| wtf21-Y-dw | TCATCTCTACCTTGATTTGAC |
| wtf22-U-up | CTATAACAAACCATCGTTCTG |
| wtf22-U-dw | ACGAGGCAAGCTAAACAGATTTGCGTTATTAATGTAGTTGTC |
| wtf22-D-up | GCTGTCGATTCGATACTAACGCTTCATTTCATCAAGATCATTC |
| wtf22-D-dw | CAAAGCTACAAACGCACTTC |
| wtf22-Y-up | AGAAGCAAGTGGTACAAATAC |
| wtf22-Y-dw | CATCAAGCTTCATATACCCC |
| wtf23-U-up | CCCATTTACACGGTATTGTG |
| wtf23-U-dw | ACGAGGCAAGCTAAACAGATTGACACATTTATTTTGTCACTG |
| wtf23-D-up | GCTGTCGATTCGATACTAACGAATGTTATAGTAGCGTGTCAG |
| wtf23-D-dw | CGTTCCTCAGTTCAGTTATG |
| wtf23-Y-up | GAGAATGTAGCATATACGTTG |
| wtf23-Y-dw | CGGTTTCTACGTTATACACC |
| wtf24-U-up | CTCTGTTGCTACTTATACAAC |
| wtf24-U-dw | ACGAGGCAAGCTAAACAGATTCTTCATAATAATAATATCATTCC |
| wtf24-D-up | GCTGTCGATTCGATACTAACGGCGTTATTAATGTAGTTGTCG |
| wtf24-D-dw | CTATTGCAATGTGTCTTTAGG |
| wtf24-Y-up | GCGATCTACTATATGATGGC |
| wtf24-Y-dw | GTACGATATCTGTAATTTGGG |
| wtf25-U-up | CAACAACGCTTACAGATAGC |
| wtf25-U-dw | ACGAGGCAAGCTAAACAGATCGATTGAACCGTTCTCTTAG |
| wtf25-D-up | GCTGTCGATTCGATACTAACGCATTTATTCTGTCACTGCCC |
| wtf25-D-dw | CATCTGTCTAATACCATCGC |
| wtf25-Y-up | CATTGATCGTGCACAAGTTG |
| wtf25-Y-dw | GGGAAGCTTCTTCAAGATAG |

**Supplementary file 1g. Primer sequences used for *wtf* construction**

| Primer name | Sequence (5' to 3') |
|---|---|
| wtf23-antidate-up | TCAGGAACATCGTATGGGTATTAAACTTCGCCTTCGACATC |
| wtf23-antidate-dw | CttcATTCCTGACAACATCTTTATGAAGAATAAATATTACCCCTTG |
| wtf23-posion-up | aaaccccggatccaagcttATGGGAGCTAACAACCCTAAC |
| wtf23-posion-dw | CATCGTCGTCCTTGTAGTCTTAAACTTCGCCTTCGACATC |
| wtf18-poision-up | aaaccccggatccaagcttATGGACATTTCGAAACTTGCT |
| wtf18-poision-dw | CATCGTCGTCCTTGTAGTCTTAGACTTCGCTTTCGGCC |
| wtf18-antidote-up | TCAGGAACATCGTATGGGTATTAGACTTCGCTTTCGGCC |
| wtf18-antidote-dw | CttcATTCCTGACAACATCTTTATGAAGAATAATTACACTTCCTTG |
| wtfcz1-up | CTAATGATTTTTATTTCTTTATGGTGCTTCTTTGAAACTTGGACC |
| wtfcz1-dw | AGTGTCCAAATTGGAATACAAACAGATGATACAATTAAAAGTAAAAC |
| wtfcz2-up | GTTTTACTTTTAATTGTATCATCTGTTTGTATTCCAATTTGGACACT |
| wtfcz2-dw | AGTGTCCAAATTGGAATACAAACAGATGATACAATTAAAAGTAAAAC |
| wtfcz3-up | GTTCTTTTTCTTTGCGAAATGGAATGTCCAGGTGCTCTCAAAAG |
| wtfcz3-dw | CTTTTGAGAGCACCTGGACATTCCATTTCGCAAAGAAAAAGAAC |
| wtfcz4-up | GAATGGTATAGCTTTTATTTTAGAAGGTATAGGAAATGCATTTG |
| wtfcz4-dw | CAAATGCATTTCCTATACCTTCTAAAATAAAAGCTATACCATTC |
